# Capillary Network Generation Framework for Estimating Volumetric Capillary Density from Histological Vascular Measurements

**DOI:** 10.64898/2026.07.10.737824

**Authors:** Zachary Harbin, Carla Fisher, Rachel A. Morrison, Hector Gomez, Sherry Voytik-Harbin, Adrian Buganza Tepole

## Abstract

Angiogenesis drives the formation and remodeling of capillary networks throughout tissue repair, regulating the vascular environment that supports healing and tissue remodeling. Experimental characterization of these processes is commonly performed using CD31-stained histological tissue sections to quantify capillary surface density and morphology throughout healing. However, these measurements provide only two-dimensional characterization of an underlying three-dimensional (3D) vascular network, limiting direct estimation of volumetric capillary density and vascular architecture. To address this limitation, an experimentally informed framework was developed to generate representative 3D capillary networks, enabling estimation of volumetric capillary density from histologically quantified vascular measurements. CD31-stained histological sections obtained from a longitudinal porcine lumpectomy study were analyzed to quantify the percentage of CD31-positive area (%CD31+) and capillary morphology within healthy tissue and healing surgical cavities. Histologically quantified morphology distributions and literature-informed vascular branching characteristics were incorporated into a capillary network generation framework to construct representative 3D vascular networks. Capillary branches were iteratively generated within representative tissue volumes until virtual histological sections reproduced experimental %CD31+ measurements, enabling estimation of volumetric capillary density. Generated capillary networks demonstrated good agreement with experimentally characterized 3D vascular architecture, while simulated histological sections accurately reproduced experimentally quantified capillary counts and vascularization measurements. Application of the framework to the porcine lumpectomy dataset captured temporal changes in vascular remodeling throughout healing, revealing progressive increases in volumetric capillary density and vascular maturation. Collectively, this framework provides an experimentally informed methodology for relating histological vascular measurements to volumetric capillary density estimates, supporting future computational studies of angiogenesis and tissue repair.

## 1 Introduction

Angiogenesis and vascular remodeling are central processes throughout tissue healing. Following tissue injury, disruption of the native vasculature and reduced oxygen availability create an inflammatory and hypoxic environment within a provisional matrix [1,2]. These signals promote secretion of cytokines and angiogenic factors that stimulate endothelial activation, angiogenic sprouting, and vascular ingrowth into the damaged region [3,4]. The developing capillary network subsequently supports oxygen and nutrient delivery needed for new extracellular matrix (ECM) deposition and remodeling [3,5]. As the inflammatory response subsides, the initially dense and highly angiogenic vascular network gradually undergoes maturation and pruning [4]. Characterizing the temporal evolution of vascular architecture throughout healing is essential for understanding tissue repair and informing computational models of angiogenesis and tissue regeneration.

Computational modeling frameworks have increasingly been developed to better understand the coupled mechanobiological processes governing wound healing and tissue remodeling. Historically, the majority of these efforts have focused on cutaneous wound healing, with a subset of models incorporating vascular remodeling and angiogenesis processes [6–16]. Despite these advances, many existing angiogenesis models remain limited by a lack of direct calibration to experimentally quantified vascular remodeling behavior, often relying on assumed parameterization and idealized healing responses.

Prior work from our group established a computational mechanobiological model for cavity healing following breast-conserving surgery (BCS; lumpectomy) [17,18]. The original model incorporated inflammatory signaling together with fibroblast activity and collagen remodeling, with model calibration informed by histopathological tissue data from a preclinical porcine lumpectomy study [17,18]. However, angiogenesis and vascular remodeling were not explicitly represented within the model formulation. More recently, ongoing work has focused on incorporating these processes to better capture the evolving biochemical and vascular environment associated with healing following BCS. These efforts leverage the same preclinical porcine lumpectomy dataset, where vascular remodeling throughout healing is characterized through CD31 immunohistochemical analysis of histological tissue sections, providing experimental measurements used to calibrate the model. However, these measurements inherently provide a two-dimensional (2D) characterization of an underlying three-dimensional (3D) vascular network, limiting direct estimation of volumetric capillary density and architecture needed to inform 3D computational representations of vascular remodeling. As a result, translating histological vascular measurements into representative 3D capillary density estimates remains a largely unresolved challenge for computational modeling of angiogenesis and vascular remodeling.

In this study, we developed an experimentally informed capillary network generation framework for translating quantified CD31-stained histological measurements into representative volumetric capillary density estimates. CD31-stained tissue sections obtained from a preclinical porcine lumpectomy study were analyzed to quantify capillary density across healthy and post-surgical healing conditions. These experimentally quantified measurements were subsequently used to guide generation of capillary networks within representative volume elements (RVEs), where stochastic branching and network formation were iteratively performed until virtual histological sections matched experimentally observed vascularization. The methodology was evaluated through comparison with previously reported vascular architectural characteristics, together with comparison between experimentally quantified and virtually estimated capillary counts from corresponding tissue sections. Collectively, this work establishes a framework for translating histological vascular measurements into representative 3D capillary density estimates, enabling experimentally informed characterization of capillary evolution and vascular remodeling throughout healing while supporting integration of vascularization data into computational angiogenesis models.

## 2 Methods

### 2.1 Histological Data Processing and Quantification

Histological sections were derived from a longitudinal porcine simulated lumpectomy study involving female Yucatan mini-pigs (45–65 kg) conducted under a protocol approved by the Purdue University Institutional Animal Care and Use Committee [2]. All animal handling and care were performed in accordance with relevant NIH and AAALAC guidelines. Simulated lumpectomies involved excision of approximately one quarter of the mammary tissue. Titanium marker clips (Ethicon Small LIGACLIP, West CMR, Clearwater, FL) were placed to facilitate margin identification of surgical sites. Explanted breast tissue was collected from healthy tissue and at 1, 4, and 16 weeks following surgery, fixed in 10% neutral-buffered formalin, and processed for paraffin embedding. Transverse sections (4 *μm*) were obtained from regions encompassing the surgical site and adjacent native tissue. Sections were immunostained for CD31 to identify endothelial-lined vascular structures. Immunolabeling was performed on deparaffinized, antigen-retrieved sections using rabbit monoclonal anti-CD31 (Abcam, Waltham, MA) and a horse anti-rabbit ImmPRESS-AP secondary antibody (Vector Laboratories, Newark, CA). Slides were counterstained with hematoxylin and scanned at high resolution using the Aperio VERSA 8 whole-slide scanner (Leica Biosystems, Deer Park, IL) under standardized imaging conditions.

CD31-stained histological sections from healthy breast tissue and 1, 4, and 16 week post-surgical cavity regions were analyzed to characterize vascular morphology across time. For each time-point, 2-3 histological sections were analyzed, with 25 representative cross-sections (500 × 500 *μm*^2^) sampled throughout the tissue domain. Regions of interest were extracted using Aperio ImageScope (Leica Biosystems, Vista, CA) and subsequently processed in ImageJ (National Institutes of Health, Bethesda, MD) to isolate and quantify CD31-positive vascular structures. Image processing consisted primarily of color-based thresholding to segment positively stained regions, followed by morphological filtering and particle analysis to remove noise. The resulting segmented images were used to compute the percentage of CD31-positive area (%CD31+), representing the fraction of the 2D histological region occupied by capillaries. A detailed description of the image processing and segmentation methodology is provided in Appendix A. In addition to %CD31+ quantification, capillaries with visible lumens were identified from a subset of segmented histological images for characterization of capillary cross-sectional geometry. Geometric features including minimum Feret diameter and capillary wall thickness were evaluated across multiple cross-sections to characterize time-dependent variations in capillary structure.

### 2.2 Capillary Network Generation

To estimate volumetric capillary density from experimentally quantified %CD31+ measurements, representative 3D capillary networks were generated using the framework outlined in Fig. 1. Histological %CD31+ measurements corresponding to the evaluated tissue timepoint, along with the RVE geometry, were specified as inputs for the capillary network generation process. For the primary simulations performed in this study, capillary networks were constructed within cubic RVEs corresponding to the dimensions of the histological cross-sections (500 × 500 × 500 *μm*^3^). Within the prescribed RVE, capillary networks were constructed through iterative generation of capillary branches informed by experimentally derived geometric and architectural characteristics. For each capillary branch, a branch order (*k*) was first assigned by sampling from branching order distributions derived from previously reported 3D micro-computed tomography (microCT) analyses of wound vascular architecture [19]. Following assignment of the branch order, an initial branch segment (*k* = 1) was generated through random assignment of a spatial node location and branch direction within the RVE. Branch lengths were independently sampled from experimentally derived distributions reported in prior 3D microCT characterization of wound microvasculature [20]. Capillary cross-sectional geometry was independently assigned for each generated branch through sampling of minimum Feret diameter and capillary wall thickness distributions quantified from the analyzed histological sections corresponding to the specified tissue timepoint. For branch structures with prescribed orders greater than one (*k* > 1), additional branch segments were sequentially generated from the existing branch structure until the assigned order was satisfied. These subsequent branch segments were generated using the same stochastic sampling procedures for branch length and capillary cross-sectional geometry, while branch directions were randomly assigned subject to geometric constraints preventing growth toward the parent branch orientation, thereby promoting outward extension of the developing capillary structure. For higher-order branch structures (*k* ≥ 3), probabilistic junction formation rules were implemented to promote physiologically representative branching behavior within the developing capillary network. Specifically, at least one branch junction, defined as a node connecting multiple branch segments, was enforced within each higher-order branch structure. The remaining nodes within the developing branch structure were then independently assigned a 50% probability of junction formation, allowing generation of variable branching morphologies throughout the simulated capillary network. Once the assigned order and associated branching structure were satisfied, the completed capillary branch was incorporated into the developing capillary network.

**Fig. 1.**
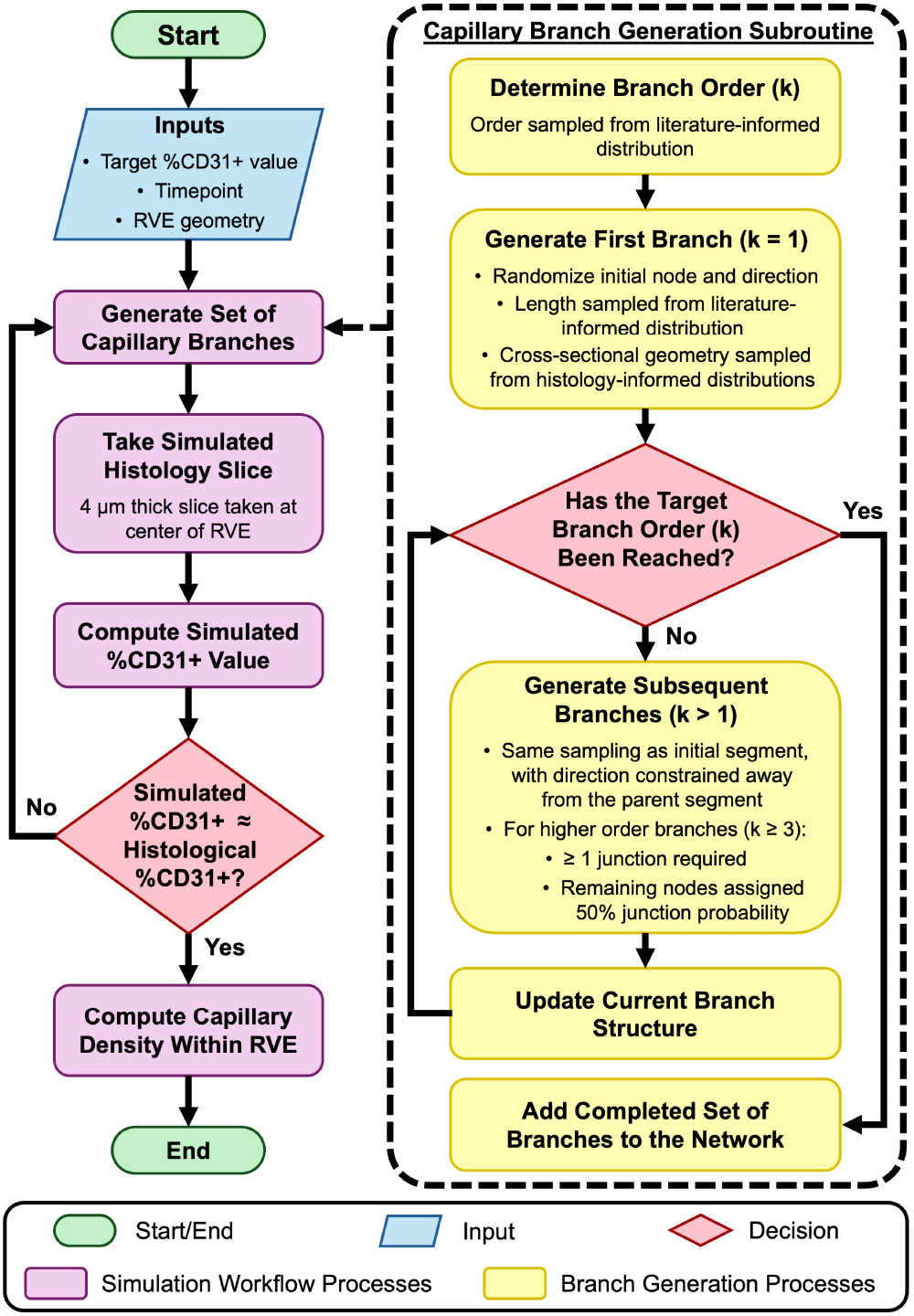
Workflow for generating 3D capillary networks and estimating volumetric capillary density from histological %CD31+ values. Associated RVE geometry and histological %CD31+ measurements obtained at the analyzed timepoint are used as inputs to the simulation framework. A set of capillary branches is then generated within the RVE, where the branch structure and order are informed through literature-derived distributions, while the capillary cross-sectional geometry is sampled from time-dependent histological distributions. Following generation of the capillary branch set, a simulated histology slice is extracted from the RVE and the corresponding %CD31+ value is computed within the slice. Capillary branch generation is repeated until the simulated %CD31+ value approximates the target histological %CD31+ measurement, after which volumetric capillary density is calculated by computing the total number of capillaries over the RVE volume.

Following incorporation of generated capillary branches into the developing network, a virtual histological slice was extracted from the simulated capillary network to enable direct comparison between simulated and experimentally measured %CD31+ values (Fig. 1). The virtual histological slice was generated by defining a fixed planar section through the center of the RVE. Slice thickness was prescribed to match the experimentally analyzed histological sections (4 *μm*). Capillary branch segments within the simulated network were evaluated for intersection with the virtual histological slice volume, including both complete and partial intersections with the slice domain. The portion of each intersecting segment contained within the slice was calculated and summed to determine the total capillary volume represented within the section. Simulated %CD31+ values were computed as the ratio of total capillary volume contained within the virtual histological slice to the total slice volume and subsequently compared against the experimentally measured histological %CD31+ value prescribed as the target input for the capillary network generation process. If the simulated %CD31+ value did not reproduce the experimentally measured histological %CD31+ value within a 1% relative tolerance of the target measurement, additional capillary branches were sequentially generated and incorporated into the developing network until the simulated %CD31+ value satisfied the prescribed convergence criterion. Volumetric capillary density was then computed as the total number of capillary branches contained within the simulated network divided by the corresponding RVE volume. To account for stochastic variability associated with capillary network generation, 50 independent simulations were performed for each analyzed condition, with volumetric capillary density reported as the mean ± standard deviation (SD) across the generated capillary networks.

## 3 Results

### 3.1 Histological Characterization of Capillary Cross-Sectional Geometry

For a subset of the analyzed histological regions of interest, capillary cross-sectional geometry was quantified to characterize vessel morphology across healthy and post-surgical tissue conditions, as shown in Fig. 2. Healthy breast tissue (Fig. 2A) exhibited a minimum Feret diameter distribution (Fig. 2A(i)) strongly skewed toward smaller capillary cross-sectional dimensions, indicating a predominance of small capillaries with relatively few larger vessels present. The average minimum Feret diameter for healthy breast tissue was 12.59 ± 7.55 *μm*. In contrast, capillary wall thickness distributions (Fig. 2A(ii)) exhibited a broader and more evenly distributed range of values, with an average wall thickness of 2.76 ± 0.97 *μm*, indicating greater variability in wall thickness across the healthy capillary population. Due to formation of a prominent seroma and/or hematoma within the cavity region at 1 week post-surgery, no identifiable capillary structures were present for geometric characterization, and therefore capillary cross-sectional analysis was not performed for this condition. At 4 weeks post-surgery (Fig. 2B), minimum Feret diameter distributions (Fig. 2B(i)) shifted toward larger capillary dimensions relative to healthy breast tissue (14.62 ± 7.17 *μm*), with substantially fewer small capillary structures observed. Capillary wall thickness distributions (Fig. 2B(ii)) similarly demonstrated reduced variability relative to healthy breast tissue, with values more strongly concentrated around intermediate wall thicknesses and an average wall thickness of 2.39 ± 0.76 *μm*. By 16 weeks post-surgery (Fig. 2C), minimum Feret diameter distributions (Fig. 2C(i)) demonstrated a broader and more heterogeneous distribution relative to the 4 week post-surgical condition, with greater representation of larger vessels while maintaining a similar average minimum Feret diameter (14.58 ± 6.29 *μm*). Capillary wall thickness distributions (Fig. 2C(ii)) shifted toward larger wall thicknesses (2.70 ± 0.61 *μm*) and exhibited a more evenly distributed range of values than the 4-week condition, consistent with continued vascular remodeling and vessel maturation during the later stages of healing.

**Fig. 2.**
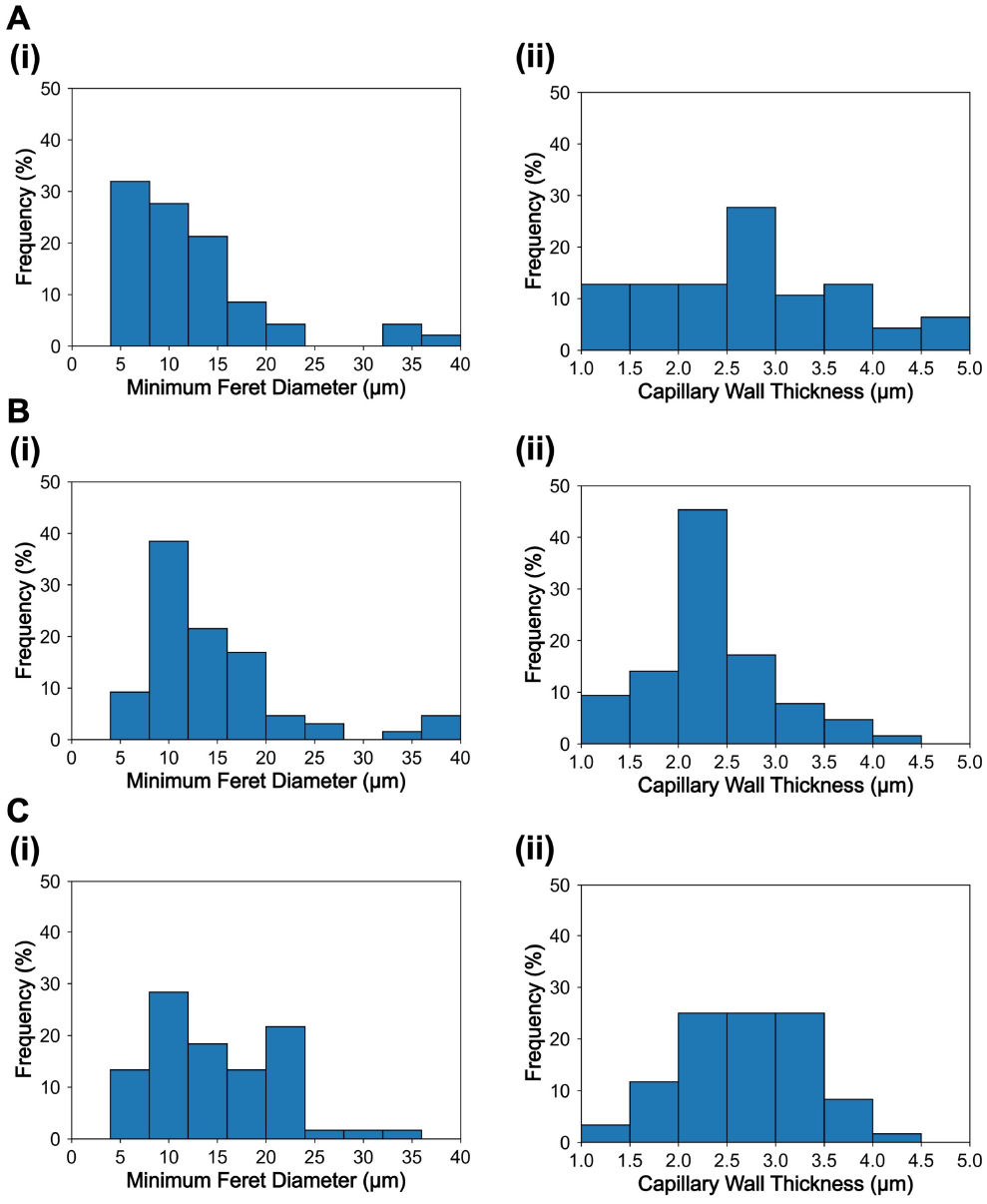
Histologically quantified capillary cross-sectional geometry distributions used to inform simulated capillary network generation. Distributions of (i) minimum Feret diameter and (ii) capillary wall thickness measured from a subset of histological capillaries with visible lumens within (A) healthy breast tissue (n = 47), (B) 4-week post-surgical cavity tissue (n = 64), and (C) 16-week post-surgical cavity tissue (n = 60).

### 3.2 Validation of Simulated Capillary Network Architecture

To evaluate the ability of the capillary network generation framework to create physiologically representative vascular architectures, we first compared simulated networks against 3D microCT measurements of cutaneous wound microvasculature reported in a previous study [20]. In contrast to the histology-based RVEs described later, validation of capillary networks against the study in [20] required a cylindrical domain matching the geometry of the experimental microCT wound analysis (8 mm diameter, 2.7 mm thickness), as shown in Fig. 3A. To enable direct comparison between simulated and experimentally quantified vascular architecture, a prescribed capillary branch count of approximately 11,660 branches was used, matching the microCT measurements [20]. A total of 50 independent networks were generated for this validation study, with a representative network shown in Fig. 3A. The resulting network architecture was then characterized by quantifying the total number of branch junctions and total capillary branch length, which were compared against the corresponding experimental microCT measurements (Fig. 3B). Simulated capillary network architectures demonstrated strong agreement with the experimental microCT measurements, with branch junction counts and total capillary branch length differing from the experimental averages by 0.88% and 8.42%, respectively, both of which remained within the reported experimental variability across skin wound samples (Fig. 3B). These results support the ability of the branching framework to reproduce physiologically representative capillary network architecture, providing a foundation for the subsequent calibration using %CD31+ histology images.

**Fig. 3.**
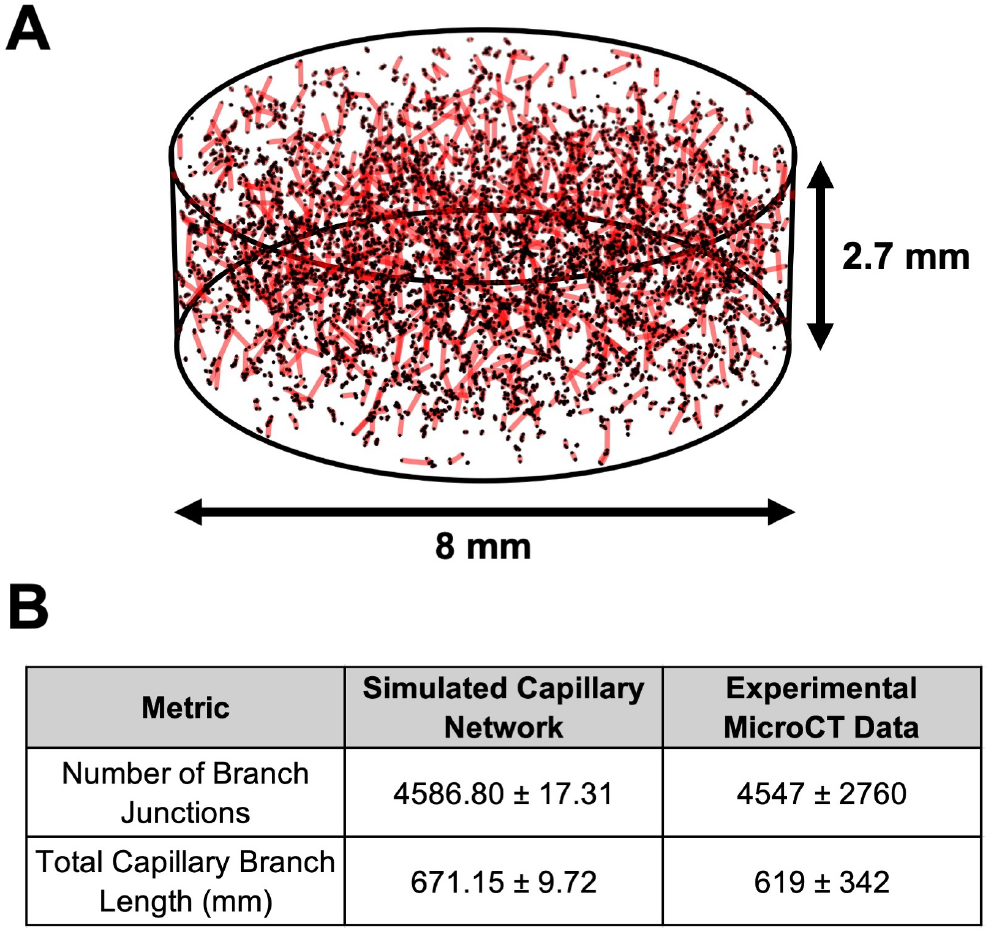
Validation of the simulated capillary network architecture against experimentally measured microCT vascular morphology. (A) Representative simulated capillary network generated within a cylindrical RVE matching the dimensions of the experimental published wound geometry used in the microCT analysis (8 mm diameter, 2.7 mm thickness) [20]. The total number of capillary branches within the simulated network was prescribed to match the experimentally reported average branch count. Red line segments represent individual capillary branch segments, while black points indicate the nodes connecting adjacent branch segments, including branch junctions and terminal endpoints. (B) Comparison of structural metrics characterizing the simulated capillary networks (n = 50) and experimental microCT vascular architecture from skin wounds (n = 6) [20], including the total number of branch junctions and total capillary branch length (mean ± SD).

### 3.3 Estimation of Volumetric Capillary Density from Histological Measurements

Representative examples of the histological %CD31+ quantification and capillary network generation process are shown in Fig. 4 across healthy breast tissue, 4 week post-surgical tissue, and 16 week post-surgical tissue conditions. Experimentally acquired CD31-stained histological sections were post-processed in ImageJ to isolate and quantify CD31-positive vascular structures, with representative segmented histological regions shown in Fig. 4A(i)-C(i). Corresponding histological %CD31+ measurements quantified across the analyzed tissue regions are summarized in Table 1 across healthy breast tissue and post-surgical healing timepoints. Healthy breast tissue demonstrated substantial differences in vascular density between adipose and fibroglandular tissue regions, where adipose tissue exhibited minimal CD31-positive staining while fibroglandular tissue demonstrated substantially greater vascularization and increased presence of capillary structures (Table 1). Due to formation of the seroma and/or hematoma within the cavity space at 1 week post-surgery, no identifiable vascular structures were present throughout the analyzed tissue regions, resulting in negligible histological %CD31+ measurements (Table 1). Increased vascularization was observed throughout the healing cavity environment at 4 weeks post-surgery, although corresponding %CD31+ values remained reduced relative to healthy fibroglandular tissue (Table 1). This vascularization continued to progress by 16 weeks post-surgery, where histological %CD31+ measurements became comparable to those observed in healthy fibroglandular tissue, consistent with progression toward a more mature remodeling state (Table 1).

**Table 1.** Histological %CD31+ measurements and corresponding volumetric capillary density estimates (mean ± SD) across healthy breast tissue and post-surgical healing timepoints.

| Tissue Type/<br>Timepoint | %CD31+ Value<br>(%) | Capillary Density<br>(cap./mm<sup>3</sup>) |
| --- | --- | --- |
| Healthy Breast Tissue |  |  |
| Adipose<br>Tissue | 0.07 ± 0.09 | 354.26 ± 435.80 |
| Fibroglandular<br>Tissue | 1.78 ± 0.82 | 5600.70 ± 2824.67 |
| Post-Surgical Tissue |  |  |
| Week 1 | 0 ± 0 | 0 ± 0 |
| Week 4 | 1.37 ± 0.37 | 3905.68 ± 1465.39 |
| Week 16 | 1.82 ± 0.67 | 4550.41 ± 1959.27 |

**Fig. 4.**
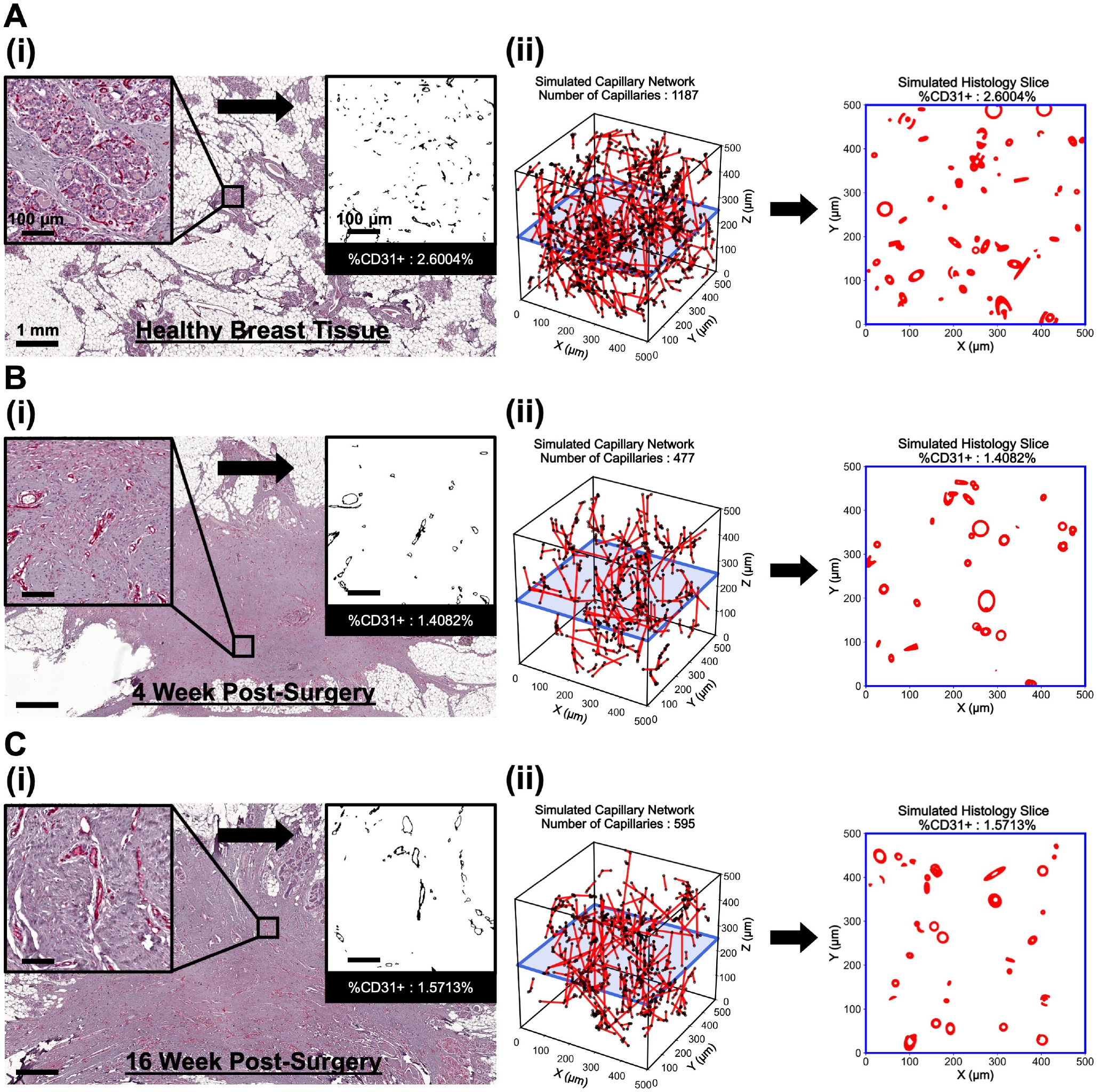
Histological quantification of capillary surface density and generation of representative simulated capillary networks for estimation of volumetric capillary density. (A) Healthy glandular tissue, (B) 4-week post-surgical cavity tissue, and (C) 16-week post-surgical cavity tissue stained with CD31 to visualize capillaries within histological sections. (i) Histological images were subdivided into 500 × 500 *µm*^2^ regions of interest to quantify the %CD31+ value representing capillary surface density. (ii) Corresponding 500 × 500 × 500 *µm*^3^ representative volume elements were iteratively generated with simulated capillary networks until simulated histology slices matched the experimentally measured %CD31+ values.

The histological %CD31+ values quantified from each individual cross-section were then used as target inputs for the capillary network generation framework to estimate corresponding volumetric capillary densities. Representative examples of the resulting simulated capillary networks are shown in Fig. 4A(ii)-C(ii). This framework enabled experimentally measured 2D histological %CD31+ values to be translated into corresponding three-dimensional volumetric capillary density estimates, with results summarized in Table 1.

To provide additional validation of the framework, we evaluated whether capillary networks generated to match experimentally measured %CD31+ values also reproduced the experimentally observed capillary counts. For a subset of histological sections, individual capillaries were manually quantified and compared with capillary counts obtained from repeated simulated histological slices generated after matching the experimental %CD31+ value. Across the analyzed histological cross-sections, all experimentally measured capillary counts fell within the approximate 95% prediction interval of the simulated capillary count distributions, and the corresponding mean simulated capillary counts differed from the experimentally measured capillary counts by an average of 14.83%. These results provide additional confidence that the framework not only reproduces the prescribed %CD31+ values, but also generates capillary architectures that are consistent with the experimentally observed vascular morphology.

## 4 Discussion

The present study provides experimentally quantified characterization of vascular remodeling throughout post-surgical breast cavity healing, where temporal changes in capillary density and morphology reflected evolving angiogenic activity throughout healing. Minimal vascularization observed throughout the 1 week post-surgical cavity environment corresponded with the formation of a seroma and/or hematoma within the cavity space, which was observed both grossly and histologically in the associated preclinical porcine lumpectomy study [2]. These findings are consistent with the early healing environment following surgery, where inflammatory cells and early angiogenic activity primarily localize along the cavity boundary while the cavity interior remains dominated by fluid accumulation with minimal cellular infiltration and capillary presence [2]. In contrast, increased vascularization observed at 4 and 16 weeks post-surgery, reflected by corresponding increases in histological %CD31+ measurements and estimated volumetric capillary density (Table 1), was consistent with progression through the proliferative and remodeling phases of healing. These observations are consistent with the established role of angiogenesis during proliferative healing, where expansion of the vascular network supports formation of fibrovascular tissue and facilitates the cellular and extracellular matrix remodeling [3]. Temporal changes in capillary morphology, including greater representation of larger vessel diameters and thicker capillary walls during later healing stages (Fig. 2), further suggest continued vascular maturation. Such changes are consistent with stabilization and remodeling of the initially immature neovasculature as the tissue transitions from active repair toward long-term remodeling [4,21]. Previous studies of cutaneous wound healing have similarly reported transient increases in vascular density during the proliferative phase followed by gradual vascular regression during later remodeling stages [20,22]. The sustained vascular density observed between the analyzed 4 and 16 week post-surgical timepoints may reflect the distinct healing environment associated with large volumetric breast cavity repair compared to cutaneous wound healing.

Experimental characterization of angiogenesis and vascular remodeling is most commonly performed through CD31 immunohistochemical staining of 2D tissue sections. However, these measurements inherently provide only planar characterization of an underlying 3D vascular network, making direct estimation of volumetric capillary density and vascular architecture nontrivial. Although additional techniques capable of characterizing 3D vascular architecture, including microCT, magnetic resonance angiography, intravital microscopy, and tissue-clearing approaches combined with optical imaging, have been utilized, these methods remain substantially less widespread and often require specialized inSTRUMENTATION and sample preparation relative to conventional histological characterization methods [19,20,23–25]. Consequently, translating routinely acquired histological measurements into representative volumetric vascular descriptions remains an important challenge. The present framework addressed this gap through stochastic generation of capillary networks informed by experimentally quantified %CD31+ measurements and capillary morphology distributions based on a thorough review of the literature. Through iterative virtual histological sectioning and matching of simulated and experimental %CD31+ values (Fig. 4), the framework generated representative volumetric vascular networks consistent with the observed histological slides. Importantly, this approach does not attempt to reconstruct the exact underlying vascular topology from individual histological sections, but rather generates representative volumetric network structures constrained by experimentally observed histology slides.

Estimation of representative 3D vascular networks from conventional histological measurements requires that the generated capillary architecture remain consistent with experimentally observed vascular morphology. To achieve this, the proposed framework incorporated experimentally quantified capillary cross-sectional geometries together with literature-derived 3D branching characteristics to constrain stochastic capillary network generation [19,20]. Independent validation against previously reported 3D microCT vascular architecture measurements demonstrated that the generated capillary networks accurately reproduced key structural characteristics of physiological wound vasculature (Fig. 3). Comparison between experimentally quantified and simulated capillary counts further demonstrated that capillary networks generated to match histological %CD31+ measurements remained consistent with the corresponding histological observations. Together, these validation studies establish confidence that the generated capillary networks provide representative volumetric descriptions of the underlying vasculature, enabling direct integration of experimentally quantified vascular remodeling into the computational mechanobiological model. This experimentally informed framework provides a foundation for future computational investigations of angiogenesis and vascular remodeling throughout tissue healing.

This study is not without limitations. While the proposed framework incorporated experimentally quantified %CD31+ measurements and capillary morphology distributions to generate representative volumetric vascular networks, additional architectural characteristics including vessel tortuosity and branching angles were not explicitly incorporated into the current formulation. Similarly, capillary morphology was characterized using minimum Feret diameter and wall thickness distributions obtained from histological cross-sections. Although these descriptors were experimentally informed, they do not fully capture the complexity of vessel morphology. Furthermore, the framework does not seek to reconstruct the exact underlying 3D vascular topology, but rather generates representative volumetric vascular architectures constrained by experimentally quantified histological measures. Consequently, multiple vascular network configurations may satisfy similar histological measurements despite differences in their underlying 3D organization. Accordingly, multiple stochastic network realizations were performed for each tissue condition to characterize the range of vascular architectures consistent with the experimental measurements. Ultimately, the proposed methodology provides an experimentally informed approach for relating conventional histological vascular measurements to representative volumetric capillary network descriptions and estimates of capillary density.

## 5 Conclusion

This study experimentally characterized vascular remodeling throughout post-surgical breast cavity healing and developed an experimentally informed framework for translating histological vascular measurements into representative volumetric capillary network descriptions. Histological analysis demonstrated progressive changes in capillary density and morphology throughout healing, reflecting continued angiogenic activity and vascular remodeling during tissue repair. By enabling direct integration of experimentally quantified vascular remodeling into the computational mechanobiological model, the proposed framework establishes a quantitative link between conventional histological characterization and computational simulation of angiogenesis. Overall, the proposed framework establishes an experimentally informed approach for translating conventional histological vascular measurements into representative volumetric vascular network descriptions, supporting future computational studies of angiogenesis and tissue healing.

## Author Contributions

Conceptualization, Z.H., H.G., S.V.-H., and A.B.T.; methodology, Z.H., R.A.M., H.G., S.V.-H., and A.B.T.; investigation, Z.H., H.G., S.V.-H., and A.B.T.; writing–original draft, Z.H., H.G., S.V.-H., and A.B.T.; writing–review & editing, Z.H., C.F., R.A.M., H.G., S.V.-H., and A.B.T.; funding and resources, Z.H., C.F., R.A.M., S.V.-H., and A.B.T.; supervision, C.F., H.G., S.V.-H., and A.B.T.

## Acknowledgment

The authors thank the Purdue University Histology Research Laboratory, a core facility of the NIH-funded Indiana Clinical and Translational Science Institute, and its staff for their assistance with histological slide preparation.

## Funding Data

Not applicable

## Data Availability Statement

The capillary network generation framework code is available in the following repository: https://github.com/zharbin/2026_CapillaryNetwork

## Appendix A

CD31 Histological Quantification Procedure

1. Identify CD31+ Vascular Structures
  a. Apply color thresholding to isolate positively stained vascular regions
    i. Hue ([Minimum, Maximum]): [190,255]
    ii. Saturation ([Minimum, Maximum]): [190,255]
    iii. Brightness ([Minimum, Maximum]): [0,255]
  b. Apply color balance adjustments
    i. Red ([Minimum, Maximum]): [0,0]
    ii. Green ([Minimum, Maximum]): [0,0]
    iii. Blue ([Minimum, Maximum]): [0,0]
  c. Convert image type from RGB color to 32-bit grayscale
  d. Apply intensity thresholding
    i. Intensity Range ([Minimum, Maximum]): [0,150]
2. Quantify CD31-Positive Vascular Structures
  a. Enable Feret diameter under “Set Measurements”
  b. Perform particle analysis
    i. Particle Size ([Minimum, Maximum]): [5,∞]
    ii. Circularity ([Minimum, Maximum]): [0,1]
  c. Retain segmented vascular structures with minimum Feret diameters between 4 and 40 *μ*m
  d. Exclude segmented structures outside the specified Feret diameter range
3. Calculate Histological %CD31+:

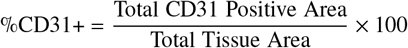
4. Characterize Capillary Cross-Sectional Morphology (Subset Analysis)
  a. Identify capillaries with visible lumens within the segmented histological images
  b. Quantify minimum Feret diameter from particle analysis measurements
  c. Estimate capillary wall thickness using local thickness analysis applied to the segmented vascular structures

## References

[1] Reinke, J. and Sorg, H., 2012, “Wound repair and regeneration,” European surgical research, 49(1), pp. 35–43.

[2] Puls, T. J., Fisher, C. S., Cox, A., Plantenga, J. M., McBride, E. L., Anderson, J. L., Goergen, C. J., Bible, M., Moller, T., and Voytik-Harbin, S. L., 2021, “Regenerative tissue filler for breast conserving surgery and other soft tissue restoration and reconstruction needs,” Scientific Reports, 11(1), p. 2711.

[3] Eming, S. A., Brachvogel, B., Odorisio, T., and Koch, M., 2007, “Regulation of angiogenesis: wound healing as a model,” Progress in histochemistry and cytochemistry, 42(3), pp. 115–170.

[4] DiPietro, L. A., 2016, “Angiogenesis and wound repair: when enough is enough,” Journal of Leucocyte Biology, 100(5), pp. 979–984.

[5] Eming, S. A., Martin, P., and Tomic-Canic, M., 2014, “Wound repair and regeneration: mechanisms, signaling, and translation,” Science translational medicine, 6(265), pp. 265sr6–265sr6.

[6] Pettet, G., Byrne, H., McElwain, D., and Norbury, J., 1996, “A model of wound-healing angiogenesis in soft tissue,” Mathematical biosciences, 136(1), pp. 35–63.

[7] Olsen, L., Sherratt, J. A., Maini, P. K., and Arnold, F., 1997, “A mathematical model for the capillary endothelial cell-extracellular matrix interactions in wound-healing angiogenesis,” Mathematical Medicine and Biology: A Journal of the IMA, 14(4), pp. 261–281.

[8] Byrne, H., Chaplain, M., Evans, D., and Hopkinson, I., 2000, “Mathematical modelling of angiogenesis in wound healing: comparison of theory and experiment,” Computational and Mathematical Methods in Medicine, 2(3), pp. 175–197.

[9] Gaffney, E., Pugh, K., Maini, P., and Arnold, F., 2002, “Investigating a simple model of cutaneous wound healing angiogenesis,” Journal of mathematical biology, 45(4), pp. 337–374.

[10] Maggelakis, S. A., 2003, “A mathematical model of tissue replacement during epidermal wound healing,” Applied Mathematical Modelling, 27(3), pp. 189–196.

[11] Schugart, R. C., Friedman, A., Zhao, R., and Sen, C. K., 2008, “Wound angiogenesis as a function of tissue oxygen tension: a mathematical model,” Proceedings of the National Academy of Sciences, 105(7), pp. 2628–2633.

[12] Xue, C., Friedman, A., and Sen, C. K., 2009, “A mathematical model of ischemic cutaneous wounds,” Proceedings of the National Academy of Sciences, 106(39), pp. 16782–16787.

[13] Machado, M. J., Watson, M. G., Devlin, A. H., Chaplain, M. A., McDougall, S. R., and Mitchell, C. A., 2011, “Dynamics of angiogenesis during wound healing: a coupled in vivo and in silico study,” Microcirculation, 18(3), pp. 183–197.

[14] Valero, C., Javierre, E., García-Aznar, J., and Gómez-Benito, M., 2013, “Numerical modelling of the angiogenesis process in wound contraction,” Biomechanics and modeling in mechanobiology, 12(2), pp. 349–360.

[15] Valero, C., Javierre, E., García-Aznar, J., Gómez-Benito, M., and Menzel, A., 2015, “Modeling of anisotropic wound healing,” Journal of the Mechanics and Physics of Solids, 79, pp. 80–91.

[16] Flegg, J. A., Menon, S. N., Maini, P. K., and McElwain, D. S., 2015, “On the mathematical modeling of wound healing angiogenesis in skin as a reaction-transport process,” Frontiers in physiology, 6, p. 262.

[17] Harbin, Z., Sohutskay, D., Vanderlaan, E., Fontaine, M., Mendenhall, C., Fisher, C., Voytik-Harbin, S., and Tepole, A. B., 2023, “Computational mechanobiology model evaluating healing of postoperative cavities following breast-conserving surgery,” Computers in Biology and Medicine, 165, p. 107342.

[18] Harbin, Z., Fisher, C., Voytik-Harbin, S., and Tepole, A. B., 2026, “Computational Modeling of Patient-Specific Healing and Deformation Outcomes Following Breast-Conserving Surgery Based on MRI Data,” Annals of Biomedical Engineering, 54(2), pp. 495–513.

[19] Urao, N., Okonkwo, U. A., Fang, M. M., Zhuang, Z. W., Koh, T. J., and DiPietro, L. A., 2016, “MicroCT angiography detects vascular formation and regression in skin wound healing,” Microvascular research, 106, pp. 57–66.

[20] Okonkwo, U. A., Chen, L., Ma, D., Haywood, V. A., Barakat, M., Urao, N., and DiPietro, L. A., 2020, “Compromised angiogenesis and vascular Integrity in impaired diabetic wound healing,” PloS one, 15(4), p. e0231962.

[21] Feng, X., Tonnesen, M. G., Mousa, S. A., and Clark, R. A., 2013, “Fibrin and collagen differentially but synergistically regulate sprout angiogenesis of human dermal microvascular endothelial cells in 3-dimensional matrix,” International journal of cell biology, 2013(1), p. 231279.

[22] Michalczyk, E. R., Chen, L., Fine, D., Zhao, Y., Mascarinas, E., Grippo, P. J., and DiPietro, L. A., 2018, “Pigment epithelium-derived factor (PEDF) as a regulator of wound angiogenesis,” Scientific Reports, 8(1), p. 11142.

[23] Kim, E., Zhang, J., Hong, K., Benoit, N. E., and Pathak, A. P., 2011, “Vascular phenotyping of brain tumors using magnetic resonance microscopy (μMRI),” Journal of Cerebral Blood Flow & Metabolism, 31(7), pp. 1623–1636.

[24] Radbruch, A., Eidel, O., Wiestler, B., Paech, D., Burth, S., Kickingereder, P., Nowosielski, M., Bäumer, P., Wick, W., Schlemmer, H.-P., et al., 2014, “Quantification of tumor vessels in glioblastoma patients using time-of-flight angiography at 7 Tesla: a feasibility study,” PloS one, 9(11), p. e110727.

[25] Lee, S., Barbe, M. F., Scalia, R., and Goldfinger, L. E., 2014, “Three-dimensional reconstruction of neovasculature in solid tumors and basement membrane matrix using ex vivo X-ray microcomputed tomography,” Micro-circulation, 21(2), pp. 159–170.

